# Revisiting the social brain hypothesis: contest duration depends on loser’s brain size

**DOI:** 10.1101/300335

**Authors:** Wouter van der Bijl, Séverine D. Buechel, Alexander Kotrschal, Niclas Kolm

**Author notes:** **Corresponding author** Name: Wouter van der Bijl, Address: Department of Zoology, Stockholm University, Svante Arrhenius väg 18B, 10691 Stockholm, Sweden.

## Abstract

**Background:** Brain size is expected to evolve by a balance between cognitive benefits and energetic costs. Several influential hypotheses have suggested that large brains may be especially beneficial in social contexts. Group living and competition may pose unique cognitive challenges to individuals and favor the evolution of increased cognitive ability. Evidence comes from comparative studies on the link between social complexity and brain morphology, but the strength of empirical support has recently been challenged. In addition, the behavioral mechanisms that would link cognitive ability to sociality are rarely studied. Here we take an alternative approach and investigate experimentally how brain size can relate to the social competence of individuals within species, a problem that so far has remained unresolved. We use the unique guppy brain size selection line model system to evaluate whether large brains are advantageous by allowing individuals to better assess their performance in a social contest situation. Based on theoretical literature, we predict that contest duration should depend on the brain size of the loser, as it is the capitulation of the losing individual that ends the fight.

**Results:** First, we show that studying the movement of competitors during contests allows for precise estimation of the dominance timeline in guppies, even when overt aggression is typically one-sided and delayed. Second, we staged contests between pairs of male that had been artificially selected for large and small relative brain size, with demonstrated differences in cognitive ability. We show that dominance was established much earlier in contests with large-brained losers, whereas the brain size of the winner had no effect. Following our prediction, large-brained individuals gave up more quickly when they were going to lose.

**Conclusions:** These results suggest that large-brained individuals assess their performance in contests better and that social competence indeed can depend on brain size. Conflict resolution may therefore be an important behavioral mechanism behind macro-evolutionary patterns between sociality and brain size. Since conflict is ubiquitous among group-living animals, the possible effects of the social environment on the evolution of cognition may be more broadly applicable than previously thought.

## Background

Brain size, relative to body size, shows a fascinating amount of variation among vertebrates [1]. In the past decades, there has been great interest in how this variation could be related to the social environment in which species live. This idea has intuitive appeal, as dealing with conspecifics who themselves are cognitive actors may be challenging [2]. Several related hypotheses have been constructed around this central premise where each places a focus on different aspects of social living, such as cunning minds of social deception in the Machiavellian intelligence hypothesis [3], pair bonding in the social brain hypothesis [4] or social learning in the cultural brain hypothesis [5]. While recent publications [6–10] debate the strength of comparative evidence linking sociality and brain size in a macro-evolutionary context, a rigorous way forward is to conduct experimental studies at the micro-evolutionary level that directly test putative mechanisms and social benefits of brain size. In sum, comparative studies provide insight via large-scale patterns and have generated strong hypotheses, yet lack the ability to demonstrate causality and thus experiments are needed to investigate putative mechanisms.

Conflict is a central aspect of social behavior in most animals and competitors often incur large costs in the contest over dominance. These costs, including physical costs such as attack damage as well as time costs and vulnerability to predation, can be reduced when contests are settled quickly, which should be facilitated by fast assessment of the expected outcome of the contest [11]. Cognitive ability is thus likely important in animal contests [12]. Here we test this recently posited hypothesis [13] that enlarged brains may provide an advantage by allowing for faster assessment during contest. More specifically, based on theoretical models of animal contests [14, 15] we hypothesized that the duration of a contest should decrease with the brain size of the loser when brain size is associated with cognitive ability [13]. This follows from the notion that the conflict is settled with the decision of the loser to abandon the fight, and it is therefore the loser’s assessment that affects contest duration [16]. The assessment ability of the winner should therefore be inconsequential. While assessment mechanisms during contests can be simple [17], such as giving up when physical exertion reaches a threshold or using direct strength comparisons during physical contact, conflicts in many species can be resolved without overt aggression and physical attacks. In contests where resolution is also not aided by display behavior and ritualization, such as the roaring and parallel walks of red deer stags [18], cognitive ability is especially likely to be important in assessment. Here, we used the guppy (*Poecilia reticulata*) as a model to study the role of brain size in behavior during conflict. We first established, combining classic and novel approaches, how dominance is formed in this system, and then we compared the assessment speed between guppies with small and large brains, which differ in cognitive ability [19, 20].

## Results

### Guppy dominance is established before overt aggression

In the first experiment we hosted 16 contests between pairs of male guppies, establishing residency for both fish by acclimating them overnight in two separated sides of a tank with an opaque divider. In the morning, the divider was removed, and the males were free to interact for two hours while being recorded from an overhead camera. Long recordings were necessary, as guppies are known to show little to no aggression in the first hour of contact [21]. We kept track of the individual identities using computer vision [22] and scored all overt aggressive interactions (attacks) in these contests, defined as chases, lunges and bites. We found that the attacks within pairs were typically extremely one-sided, with on average 90% of the attacks performed by one of the two individuals. More importantly, attacks were one-sided at the onset of aggression as there was no discernable change in the identity of the aggressor over the duration of the trial, either measured as time or attack number (binomial GLMM with random intercept and slope per trial; time: z = −0.017, p = 0.987; attack number: z = −0.020, p = 0.984). This contrasts with classic theoretical concepts of competitive situations where opponents exchange blows while collecting information [14]. Instead, aggression was one-sided and invariable with time, and so assessment must have been occurring before the onset of attacks.

A dominance relation may be apparent from other behavioral cues than overt aggression. The relative movement between individuals is often heavily informed by social status, where subordinates will avoid dominant individuals but not *vice versa.* This asymmetry in displacement has a long history in being used to assess dominance in ethological studies of e.g. children [23], macaques [24] and chickens [25]. We formalized this approach by using computer vision to track the relative position and orientation of competitors and calculated a simple displacement metric, 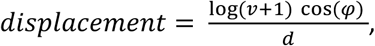 using the speed (*v*), relative orientation (*φ*) and distance (*d*) between the fish (figure 1). The displacement becomes positive and large for subjects that are quickly moving towards a competitor at short distance, and negative during a social escape.

**Figure 1:**
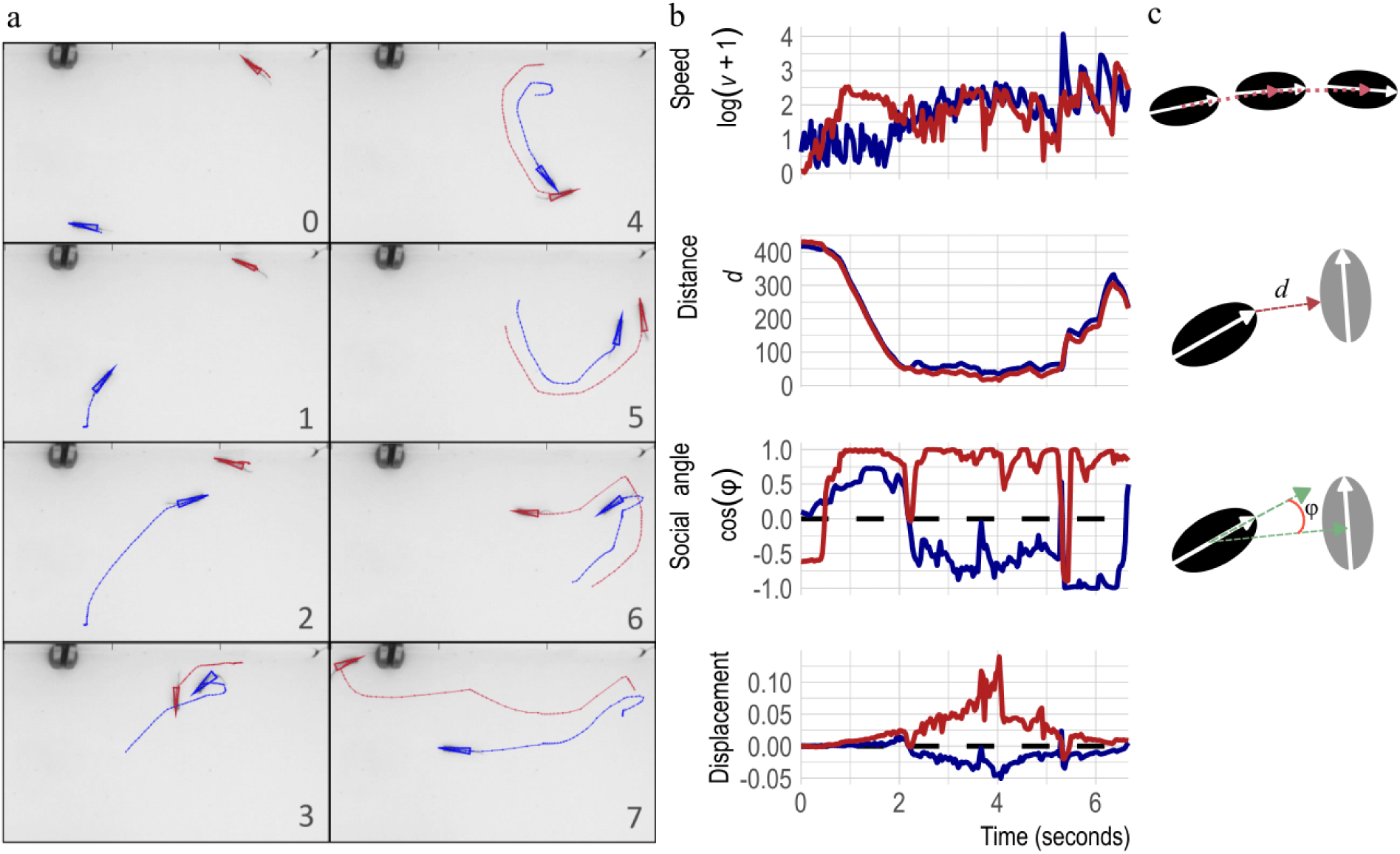
An example of quantifying male-male interactions with the displacement metric. a) Video snapshots during a contest, where the blue individual displaces the red individual, with grey numbers indicating time in seconds. Overlaid are the movement tracks used to perform the calculations. b) A timeline of 7 seconds for each of the variables and the calculated metric during the displacement event from panel a. c) Illustrations of the three variables used to calculate displacement, speed, distance and social angle.

This metric was verified to respond to discrete moments of overt aggression and distinguishes the attacker and the attacked (figures 2a & 2b). Additionally, it responds to any other, more subtle, asymmetries in movement over continuous time. That is, males spend most time interacting in close proximity and, while most of this movement is not clearly aggressive to the human observer, it is informative to the social roles as the dominant individual typically displaces the subordinate. The displacement score aims to quantify this relative movement, distinguishes between the two social roles and weighs fast and close interactions more than slow and distant ones. We confirmed that the measured displacement became asymmetrical before the onset of overt aggression by analyzing the difference in displacement scores for each trial in the time up to one minute before the first attack. In 11 out of 12 trials where attack direction clearly indicated dominance, the asymmetry in displacement before the first attack correctly predicted the eventual dominant individual (figure 2c, binomial test, p = 0.006). Therefore, the displacement scores can detect subtle movement asymmetries between opponents and correctly determine the dominant individual before classic methods could do so (figures 2d, 2e & 2f). The end of a guppy contest, and therefore the speed of assessment, is thus best described as the moment at which asymmetrical movement is established (figure 2g).

**Figure 2:**
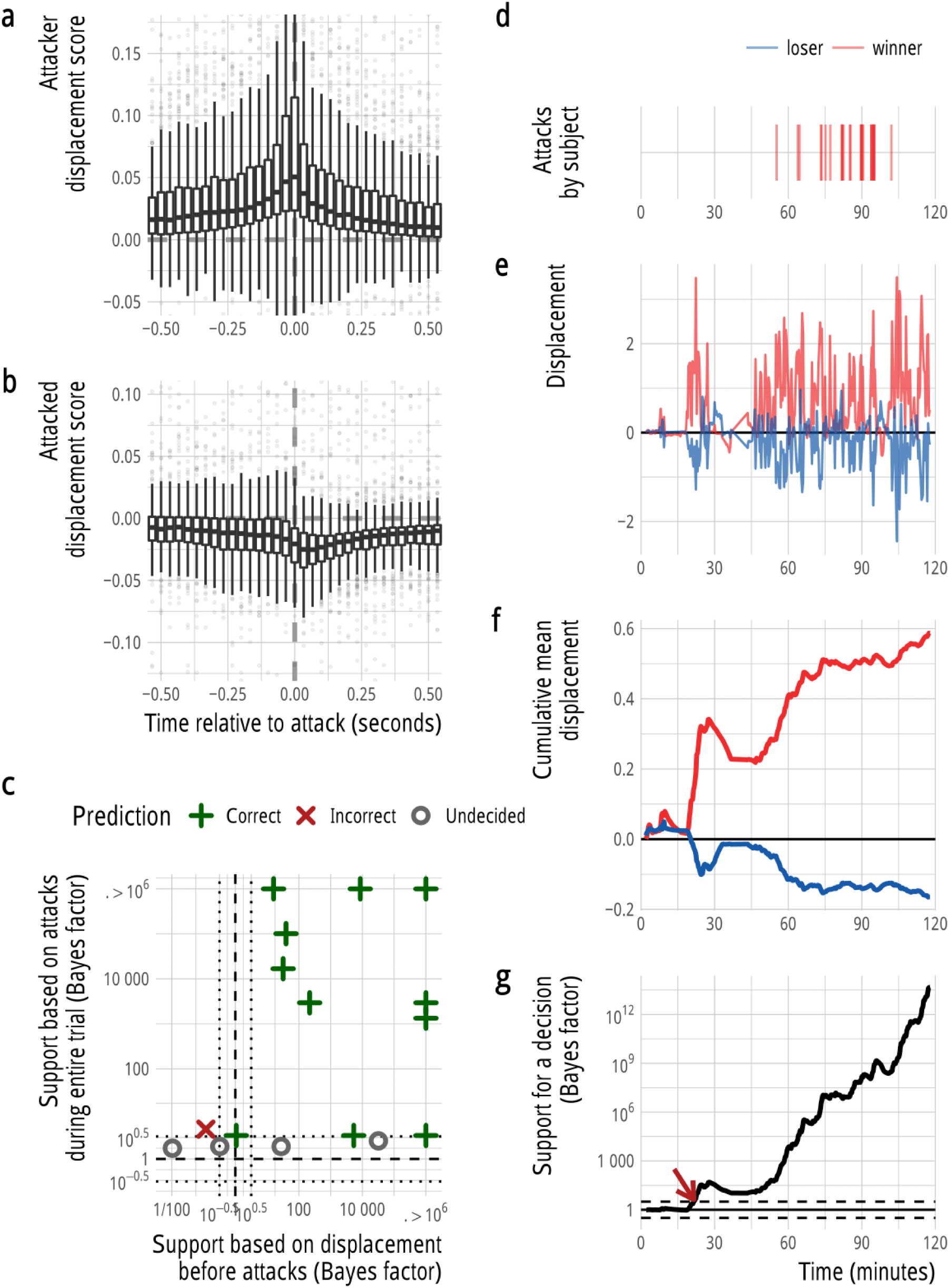
The displacement metric during overt aggression and throughout one trial. During attacks, attackers had positive displacement scores (a), while attacked fish (b) responded with negative scores, depicted by boxplots for the 1 second (30 frames) surrounding each attack. But displacement was typically already asymmetric before the first attack, since displacement preceding any attacks correctly predicted contest outcome (c). Note that Bayes Factor support larger than 10^6^ is plotted on the graph boundary. d, e, f & g depict behavior during a single trial. The timeline of attacks are shown in a), where the first attack occurred after around 50 minutes and is by one individual only. Both before and during the attacks, displacement was asymmetrical (e) and the mean displacement diverged as the trial progressed (f). Over time, the empirical support for established dominance increased as the fishes’ movements became more asymmetrical (g). When a threshold was passed (10^1/2^, dotted lines) a contest duration could be assigned (red arrow).

### Contests with large-brained losers were decided more quickly

In the second experiment, we hosted 72 contests, each lasting three hours, between male guppies from three replicated artificial selection lines for relative brain size [19]. Guppies from these selection lines have around a 10% divergence in relative brain size, with no difference in body size [19, 20]. Large-brained males from these selection lines have previously been shown to perform better in spatial mate search task [20] and to better adjust their mating preference in regards to female size [26], while large-brained females outperformed small-brained females in a numerosity task [19] and a reversal learning test [27], survived better under predation [28], responded more cautiously to a model predator [29] and displayed stronger preference for more colorful males [30]. Our experimental design mimicked a scenario of random encounters, and therefore consisted of small-brained *versus* small-brained, large-brained *versus* large-brained and small-brained *versus* large-brained, with 18, 18 and 36 contests respectively. This full factorial design allowed us to disentangle brain size effects of winners and losers. Within the contests of mixed brain size, small- and large-brained competitors were equally likely to win (19 vs 17, binomial test: 53%, p = 0.87).

We used the displacement statistic developed for the first experiment to estimate contest durations. We estimated empirical support for a difference in displacement in favor of the eventual winner across the trials. That is, at each time point in a trial we checked whether the displacement score up to that point was asymmetric between the individuals (figure 2f). To this end, we calculated Bayes factors comparing the evidence that individual 1 was winning the contest (displacement_1_ > displacement_2_) against the evidence that individual 2 was winning the contest (displacement_2_ > displacement_1_, since a classical null-hypothesis of no difference holds little biological relevance under a scenario where all pairs eventually establish dominance [21]. Once the evidence in favor of the winner outweighed that in favor of the loser by a factor 10^1/2^ (a standard cut-off point [31]) we recorded the dominance as established (figure 2g). Sixteen trials never reached this threshold and were assigned a contest duration of the maximum three hours. The estimated contest duration was not related to the total absolute displacement scores during the trial (r = −0.008, p = 0.95), and therefore long and short contest durations are unlikely to be a result of inactivity or lack of aggressive motivation, but rather the asymmetry in displacement determined our measure of contest duration.

Contests where the winner is of a much higher resource holding potential (RHP) than the loser are predicted to be shorter than those where the contestants are evenly matched or where the roles are reversed. Since previous studies do not agree on what traits predict RHP, but have indicated both size [21] and coloration [32] related traits in the guppy, we quantified standard length, tail size, condition, black coloration and bright coloration (orange, yellow and iridescence) for all males to control for such variables. Different models of assessment have different predictions in regards to how RHP is related to contest duration [16], and the trait of the loser, winner or their difference could be important. As we have no prior information on what assessment strategy guppies use and the goal of this study is to investigate brain size effects, we elected to remain agnostic to what variables should be important and used a broad model selection approach. Additionally, we included total displacement (as an estimate of total social activity) as a covariate to control for possible effects of aggressive activity. We fitted linear mixed models for different combinations of predictors, excluding models that contained the winner, loser and difference trait value of the same trait (since these models are rank deficient), and models with more than six terms (to reduce overfitting). We controlled for replicate selection line by including random intercepts for the interaction between the brain size of the winner and the brain size of the loser, nested within replicate (*lme4* syntax: (1 | replicate / (brain_size_winner : brain_size_loser)).

We found considerable model uncertainty, with 43 models that were within 4 AICc of the best model (table S1). Brain size of the loser was the most important variable and present in all models, but the inclusion of covariates varied. To take this covariate uncertainty into account we averaged coefficients over the 43 models, using full model averaging [33]. In accordance with our hypothesis, contests with large-brained losers lasted on average 36 minutes (or 34%) shorter than contests with small-brained losers (SE_adjusted_ = 13.52, z = 2.66, p = 0.008, figure 3, table S2). The effect of loser brain size was robust against making different choices in cut-off values (figure S1). The brain size of the winner, however, had no significant effect on contest duration (β = −18, SE_adjusted_ = 16.75, z = 1.072, p = 0.284, figure 3, table S2).

**Figure 3:**
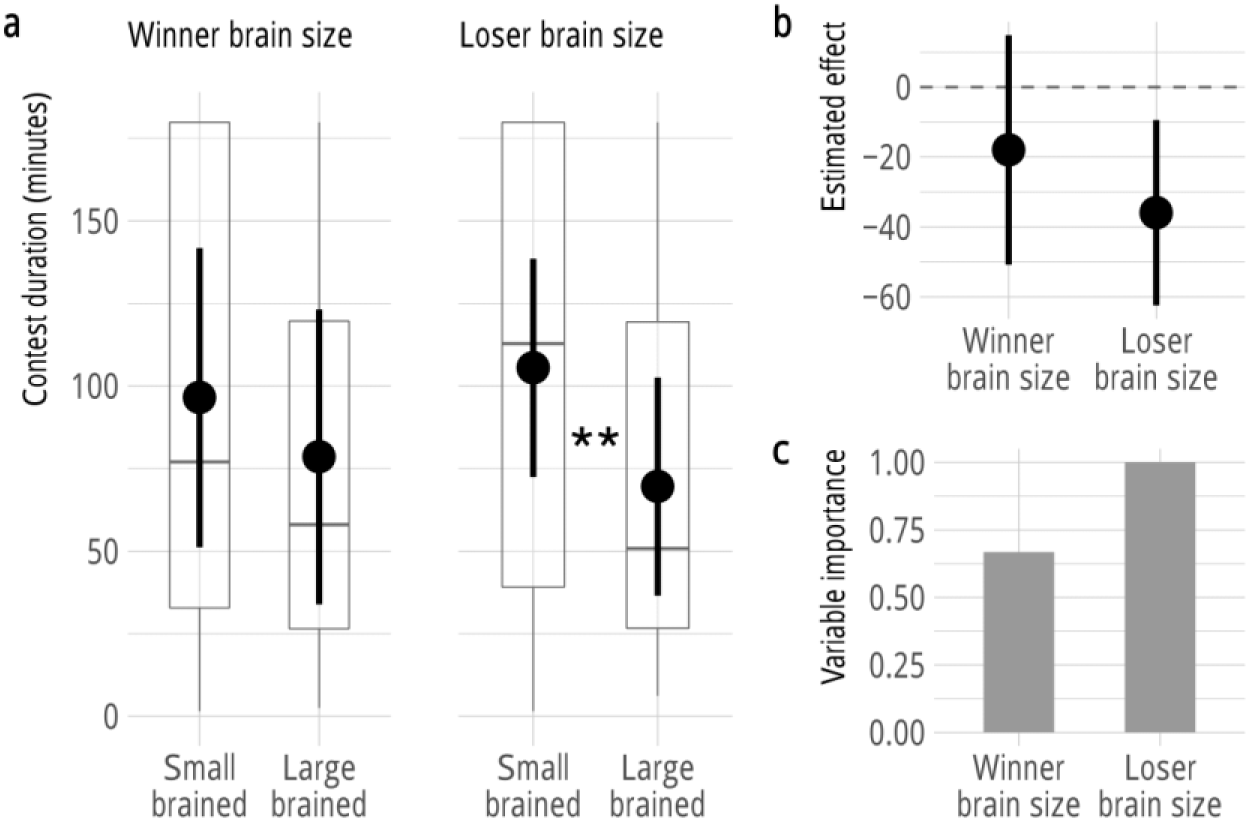
Contest duration depends on the loser’s brain size, but not the winner’s. a) Estimated contest durations as a function of brain size of either the winner, or the loser. Boxplots show the raw data, while point estimates and error-bars indicate the predicted means and 95% confidence intervals by the average model, keeping all covariates at their mean value (i.e. the marginal means). b) Estimated effect of winner and loser brain size on contest duration with 95% confidence intervals. c) Variable importance of winner and loser brain size, as the proportion of top models (within 4 AIC of the best model) that the terms are included in.

Two co-variates remained significant in the average model, the condition of the winner (β = − 21.7, SE_adjusted_ = 8.09, z = 2.68, p = 0.007, table S2) and the difference in bright coloration between the winner and the loser (β = 25.2, SE_adjusted_ = 8.57, z = 2.94, p = 0.003, table S2). When the winner was in better condition (the residual of a regression of weight on standard length), contests were shorter. This may indicate that losers assessed their opponent’s condition, or other co-varying traits. The effect of bright coloration runs in the opposite direction from expectation where more dull winners, compared to the loser, resulted in shorter contests. A possible explanation for this result is that the amount of bright coloration often changes with age, and young males are duller. This may mean that fights were shorter when winners were younger than their rivals.

## Discussion

Our results show that contest duration in the guppy depends on the brain size of the loser but not on the brain size of the winner. This demonstrates experimentally that brain size can affect the performance in a social task. Since dominance and contests are common among social animals, cognitive variation in assessment ability has the potential to be important across many taxa. Indeed, a recent study [34] showed that the amount of agonism in primate groups is positively associated with brain and neocortex size, independent of group size.

While guppies are not typically known to be a fighting species, escalated aggression and dominance hierarchies are commonly observed [21, 32]. Moreover, dominant males sire far more broods than subordinate males, independent of their attractiveness, especially when they can freely interact and jockey for position [32], indicating strong potential fitness benefits. Yet, it remains unclear to what extent overt aggression occurs in the wild, which has led some to suggest that male-male competition is perhaps not very important in this species [35]. That dominance is usually established before overt aggression, however, suggests that the absence of clearly aggressive behavior should not be interpreted as a lack of social roles since subordinate males may simply escape after dominance has become established. This means that male-male competition may play a more prominent role in the guppy than previously thought, with potentially important consequences for the study of this model system in aspects of mate choice and sexual selection.

Since contest duration could not be established by the classical method, *i.e.* simple quantification of discrete behaviors such as attacks, we developed a novel method of tracking the development of social relationships. By formalizing the well-established concept of displacement in a strongly quantitative and continuous form using modern computer vision tools, we could analyze the establishment of dominance in detail. Since in many animals overt aggression is not necessary for hierarchies to form, similar approaches could prove valuable to the broader study of contests and dominance.

Interestingly, in addition to the assessment during conflict discussed here, we have previously found guppies from the selection lines to perform better in assessment of potential mates [26, 30], as well as of the threat of predation [28, 29]. These differences were not due to differences in color perception [30] or visual acuity [36], making it unlikely that behavioral differences stem from alterations to low-level perceptual systems. Instead, volumetric increases in both the optic tectum and telencephalon [37] could have led to better integration of visual information and/or decision making in these contexts.

## Conclusion

There is an ongoing debate about macro-evolutionary correlations between measures of social complexity and brain size [8]. Till now, studies have focused on wide categorization of sociality such as social system, group size or social repertoire size, with speculations about mechanisms. Here we presented results from an experimental assessment of the link between brain size and assessment during contest, confirming our hypothesis the brain size can aid in conflict resolution. We provide a novel and complementary approach to the study of social cognitive evolution, where the experimental study of brain size effects on social cognition within species can generate bottom-up predictions to explain inter-specific variation in brain size. We hope to see further investigations into the interplay between cognition, brain size and these types of social interactions in the future. This is necessary to provide exhaustive testing of the intuitively attractive idea of a link between sociality and the brain [2, 4].

## Methods

### Experimental set-up

The first experiment used sexually mature male wildtype guppies (*Poecilia reticulata*). All individuals came from long-term lab populations whose founders were imported from Trinidad in 1998 and since then kept in large populations (>500 individuals) to avoid genetic bottlenecks.

The second experiment used fifth generation males from an artificial selection experiment on relative brain size [19]. The initial selection experiment was performed by using indirect selection; in short, random pairs were formed from the same population we used in experiment 1, and allowed to breed. After reproduction, the parents were sacrificed and their brains were weighed. The offspring from parents that had the 20% largest and 20% smallest combined relative brain size (residuals from a regression of brain weight on standard length) were then used to start the “up” and “down” selected lines. This process was repeated three times to form the three independent replicates. For a further 2 generations, the top and bottom 20% was used for further selection. The fourth and fifth generations were bred randomly to maintain the selected populations. For more details see [19].

In both experiments, individuals were housed in single-sex groups ranging from 10 to 40 individuals since early signs of sexual maturation. Fish were fed daily, alternating between flake food and freshly hatched brine shrimp for six days a week. The lab is kept at a constant 26 degrees air temperature, water temperature of 25 ± 1°C and a 12L:12D light cycle. All animals were healthy.

All experiments were performed in agreement with Swedish law based on ethical applications approved by the Stockholm Ethical Board (Reference number: N173/13, 223/15 and N8/17).

### The contests

The behavioral experimental procedure was identical in both tasks, with the exception that we recorded contests for two hours in experiment 1, and three hours in experiment 2. Contestants were chosen randomly from non-adjacent tanks, so that there had been no previous visual contact.

In the late afternoon, competitors were placed in opposite ends of an acrylic experimental tank, measuring 15×15×40 cm with 2 liters of water (3.3 cm water depth). The shallow water facilitated accurate tracking and identification. In the center of two long walls, U-shaped brackets were glued to the tank between which a wall could be tightly fit, creating two 15×20 cm compartments with no contact between the fish. The walls of the tank were covered with white plastic adhesive material and placed on a white plate to maximize contrast and aid visibility in overhead recordings. Both compartments were aerated during the night. The following morning, the aeration and wall were removed, and the fish were free to interact. After the end of the trial, the animals were separated again by placing back the wall.

All competitors were taken randomly from their home tanks. We maximized the independence of the individuals by sampling the 144 competitors from 89 difference source tanks. We randomized the brain size and replicates between days, test tanks and starting side, by running 6 batches of 12 combinations (3 replicates and four brain size combinations) and completely randomizing within each batch. All contests were between males from the same replicate selection line. We hosted 4 contests each day.

The experiment was done blindly to the brain size of competitors, as the tanks of males were marked with running numbers as soon as they were caught from their source tank. The experimenter remained blind during the moving of fish, the start and end of the trial, the measuring of competitors, the analysis of photographs, and the tracking of the videos. All contest durations were assigned algorithmically.

### Competitor traits

Directly after the trials, the competitors were photographed, their size was measured using digital calipers to the nearest 0.01 mm and weighed to the nearest 0.0001 gram. Both standard length (from the tip of the snout to the end of the caudal peduncle) and total length (from the tip of the snout to the most caudal part of the tail fin) was measured, and tail size was calculated as their difference. Both sizes and weights were measured in *duplo* and averaged. Coloration was measured by taking a photograph of the fishes left flank using a Canon EOS 1200, and then manually outlined the color patches in paint.NET and taking their area in pixels. We then calculated % body area by comparing it to the total area of the left flank in pixels. We separately quantified black, orange, yellow and iridescent coloration, but collated the latter three into a single “bright coloration” variable to reduce the number of variables in the analysis and thus avoid over-parameterization. We did not obtain pictures for two males, which reduced our sample size to 70 trials.

### Identification and tracking of animals

We started an overhead recording just before removing the opaque barrier and ended the recording after placing the barrier back. We then used idTracker [22] to keep consistent identities during the trial, and noted afterwards whether the fish had returned to their starting compartment or had been switched, in order to correctly assign the size, weight and coloration measures afterwards. In order to also obtain information about orientation and shape measurements in the tracks, we also tracked the video’s with Ctrax [38]. We matched the identities of idTracker to the detailed tracks of Ctrax by minimizing the squared errors. This allowed to also use the double tracking to automatically correct errors, by marking frames where idTracker did not corroborate the Ctrax results as missing. Leveraging the strengths of both algorithms, we obtained tracks with a median completeness of 100% and 97% in experiments 1 and 2 respectively.

### Overt aggression

We quantified overt aggression for experiment 1 by manually reviewing the two hours in each of the 16 videos and recording the time of each attack, as well as the identity of the aggressor. We scored bites and lunges, two behaviors that both are based on aggressive fast approaches but while contact was made between the individuals during a ‘bite’, no contact was observed during a ‘lunge’. Additionally, we identified chases where one individual closely and rapidly followed the other over at least several centimeters.

### Displacement

We calculated the displacement metric for both individuals separately. Speed *v* was taken as the number of pixels the centroid had moved since the last frame. Distance *d* was taken as the distance from the front of the body (the ellipsoid shape estimated by Ctrax) to the nearest point on the body of the competitor. Distance from fish *a* to *b* is therefore not necessarily the same as the distance from fish *b* to *a.* Finally, we determined the social angle *φ* as the difference in angle between the orientation of the focal fish (i.e. not the heading, but the physical orientation as estimated by Ctrax) and the angle towards the centroid of the competitor. Its cosine is therefore 1 when oriented towards the competitor, and −1 when facing away. We then combined the three variables in a displacement score as *displacement = (log(v + 1) ^∗^ cos(φ)) /d*. Speed was log transformed and we added 1 to preserve its sign. The displacement metric is strongly positive when the focal fish is quickly swimming towards the opponent at close distance, and strongly negative when it’s doing the same away from the opponent.

### Contest duration

To determine the contest duration for a trial, we calculated the displacement statistic for each individual across three hours. For each point in time we then evaluated the past displacement to check for asymmetry by calculating the Bayes Factor between two hypotheses: displacement_1_ > displacement_2_, and displacement_2_ > displacement_1_. Each of these hypotheses was evaluated using the ttestBF function from the BayesFactor R package [39], using default priors. Importantly, it is not possible to take each frame as an independent sample, as there is temporal autocorrelation. Because of this, we chose to treat every 10 seconds (300 frames) as a sample, as the autocorrelation had dropped below 0.01 at that point. The first time point where the Bayes Factor was larger than 10^1/2^ [31] was chosen as the contest duration. Our results were robust to alternate choices of the sampling interval and the Bayes Factor cut-off (figure S1).

### Statistical analysis

For details on the models used see the main text, as well as tables S1 and S2. All analyses were performed in R [40]. We fitted the (generalized) linear mixed models using lme4 [41], and performed model averaging using the MuMIn package [42]. We treated each pair of males as the statistical unit, and there were 70 trials with complete data. We visually confirmed that the residuals of the best fitting model were normally distributed.

## Declarations

### Ethics approval and consent to participate

All animal experiments were approved by the Stockholm Ethical Board (Dnr: N173/13, 223/15 and N8/17).

### Consent for publication

Not applicable.

### Availability of data and materials

All data will be publically deposited upon acceptance. R functions for loading and matching idTracker and Ctrax tracks, and the calculation of tracking statistics in R is freely available as an R package on GitHub: www.github.com/Ax3man/trackr.

## Author contributions

WvdB and NK conceptualized the study. WvdB performed the experiments, designed the methodology, performed the analyses, wrote the first draft and visualized the data. AK and NK created the selection lines. NK provided supervision and obtained funding. SDB was responsible for lab management and animal husbandry. All authors contributed to the final manuscript.

## Acknowledgements

We thank Alberto Corral López, James Herbert-Read, Björn Rogell, Karl Gotthard and Olof Leimar for fruitful discussion and helpful suggestions.

